# The mechanical microenvironment regulates ovarian cancer cell morphology, migration, and spheroid disaggregation

**DOI:** 10.1101/238311

**Authors:** Andrew J. McKenzie, Stephanie R. Hicks, Kathryn V. Svec, Hannah Naughton, Zöe L. Edmunds, Alan K. Howe

## Abstract

There is growing appreciation of the importance of the mechanical properties of the tumor microenvironment on disease progression. However, the role of extracellular matrix (ECM) stiffness and cellular mechanotransduction in epithelial ovarian cancer (EOC) is largely unknown. Here, we investigated the effect of substrate rigidity on various aspects of SKOV3 human EOC cell morphology and migration. Young’s modulus values of normal mouse peritoneum, a principal target tissue for EOC metastasis, were determined by atomic force microscopy (AFM) and hydrogels were fabricated to mimic these values. We find that cell spreading, focal adhesion formation, myosin light chain phosphorylation, and cellular traction forces all increase on stiffer matrices. Substrate rigidity also positively regulates random cell migration and, importantly, directional increases in matrix tension promote SKOV3 cell durotaxis. Matrix rigidity also promotes nuclear translocation of YAP1, an oncogenic transcription factor associated with aggressive metastatic EOC. Furthermore, disaggregation of multicellular EOC spheroids, a behavior associated with dissemination and metastasis, is enhanced by matrix stiffness through a mechanotransduction pathway involving ROCK, actomyosin contractility, and FAK. Finally, this pattern of mechanosensitivity is maintained in highly metastatic SKOV3ip.1 cells. These results establish that the mechanical properties of the tumor microenvironment may play a role in EOC metastasis.

## Introduction

Cells interpret and respond to the mechanical properties (e.g. stiffness and topology) of the extracellular matrix (ECM) by exerting contractile force and sensing counter-tension through mechanocellular systems^1^. Components of these systems include integrins, focal adhesion complexes, the actin cytoskeleton, and associated molecular motors^1,2^ serve to interpret both intrinsic and extrinsic mechanical forces into diverse signaling events such as ion flux and phosphorylation cascades. The translation of mechanical forces into biochemical signals – mechanotransduction – has been shown to regulate nearly every facet of cellular life, including shape, migration, survival, proliferation, and differentiation^3–7^. Importantly, all of the aforementioned cellular processes are altered during the pathogenesis of cancer and there is growing appreciation of the role of mechanotransduction and the mechanical microenvironment in tumorigenesis^6,8,9^. This has been particularly well-studied in the context of breast cancer, wherein tumor progression is characterized by progressive stiffening and remodeling of the tumor-associated stromal tissue and ECM^10–12^. Additionally, increases in mammographic density are associated with an increased risk for breast cancer^13,14^ and nonlinear optical imaging methods such as multiphoton microscopy (MPM) and second harmonic generation (SHG) imaging have been used to visualize local changes in collagen fibril density around invasive breast tumors^15,16^. Indeed, the increased density and reorganization of collagen fibrils around malignant breast tumors appear to facilitate local tumor cell invasion, trafficking towards blood and lymph vessels, and distal metastasis^10,15,16^. Finally, reduction of increased tumor cell-ECM tension or of matrix stiffening can normalize the malignant phenotype of primary breast cancer cells in culture and *in* vivo^10,13^.

Several observations suggest that the progression and spread of EOC may similarly be affected by ECM tension and mechanical forces. As in breast cancer, EOC and its peritoneal implants are often fibrotic and surrounded by reactive, desmoplastic stroma^17–20^, characterized by up-regulation of factors that regulate ECM content, assembly, and/or cell-matrix interactions. Also, similar to breast tumor-associated collagen signatures^15^, multi-photon and second-harmonic generation imaging has revealed altered collagen fibril density and topology associated with both primary and disseminated EOC^21–23^. Unlike breast cancer, the minimal metastatic unit of EOC is thought to be a multi-cellular aggregate, or spheroid, that exfoliates from the primary tumor and moves throughout the peritoneal cavity by normal peritoneal fluid flow^17,20,24^. These spheroids have been shown to use myosin-generated force to clear the mesothelium of peritoneal organs and attach to the submesothelial connective tissue^25^. Moreover, spheroids formed from highly-invasive EOC cells show increased expression of both lysyl oxidase 1 (LOX1), a collagen-crosslinking enzyme crucial for promoting matrix stiffness and invasion in breast cancer^10^, and tissue transglutaminase (TG2), which also promotes ECM polymerization and cross-linking^26,27^. In addition to direct ECM remodeling, inflammation associated with EOC cell implantation also alters the peritoneal/mesothelial surface and promotes a desmoplastic stromal response^28^. Moreover, there is growing consensus that endometriosis – and the associated inflammation and fibrosis in extrauterine tissues including the peritoneum and ovaries – increases the risk of EOC^29^. Increasing matrix rigidity also decreases expression of the Wnt pathway inhibitor DKK1, promoting increased Wnt signaling enhanced MT1-MMP expression and increased matrix invasion^30^. While these observations strongly implicate a role for the mechanical microenvironment in the pathogenesis of EOC, the effects of matrix rigidity on the morphological and migratory aspects of EOC cells remain largely unexplored.

In this work, we fashioned polymer hydrogels with elastic properties that mimic those of the peritoneum, a principal physiologic target for EOC dissemination, to investigate the role of substrate stiffness in EOC cell morphology and migration. We show that cell morphology, focal adhesion and cytoskeletal organization, and actomyosin contractility of SKOV-3 human EOC cells all positively correlate with substrate stiffness. Moreover, we find that matrix rigidity enhances EOC cell migration: individual EOC cells exhibit robust durotactic behavior multicellular EOC spheroids disaggregate much more efficiently on stiffer substrates. These observations establish the rigidity of the microenvironment as a potentially important facet of the pathogenesis of EOC.

## Results

### Modeling the stiffness of normal mouse peritoneum using polyacrylamide hydrogels

Little is known regarding the mechanical properties of tissues associated with the onset, progression, and spread of ovarian cancer. The peritoneum represents a major target for EOC dissemination and implantation. To measure the mechanical properties of the lining of the peritoneal cavity, we utilized atomic force microscopy (AFM) on mouse peritoneal sections (Figure 1A) to generate force-map grids (Figure 1B) and color-coded heat maps (Figure 1C) of the Young’s elastic moduli in 20 x 20 μm regions of normal mouse peritoneum. Using a custom ImageJ macro, we were able to visualize local variation of modulus values within a single force-map (Figure 1C) or global variations between numerous force maps across several different animals (Figure 1D). Global analysis revealed the average Young’s modulus of normal mouse peritoneum to be 4.33 ± 0.095 kPa (Figure 1E). While the values of the vast majority of the tissue fell within a narrow range (Figure 1E, boxed region), there were also discrete localized regions with Young’s moduli that were consistently ~10-fold higher than the average (Fig 1D *right panel*; Figure 1E, upper whisker, maximum: 35.88 kPa). These areas of stiffness ‘hotspots’ were sparse (~1 region in every 15 [20 x 20 μm] region), not coincident with any discernable underlying anatomy (e.g. blood vessels), but were consistently found in different sections take from different animals, suggesting that there is normal variation in the mechanical properties of mouse peritoneum. Polyacrylamide hydrogels were fabricated with average Young’s moduli close to the average (3.06 ± 0.03 kPa) or the maximum (25.56 ± 0.12 kPa) measured values of normal mouse peritoneum (Figure 1E), to serve as experimentally tractable yet physiologically relevant adhesive substrates to cells for subsequent experiments.

**Figure 1.**
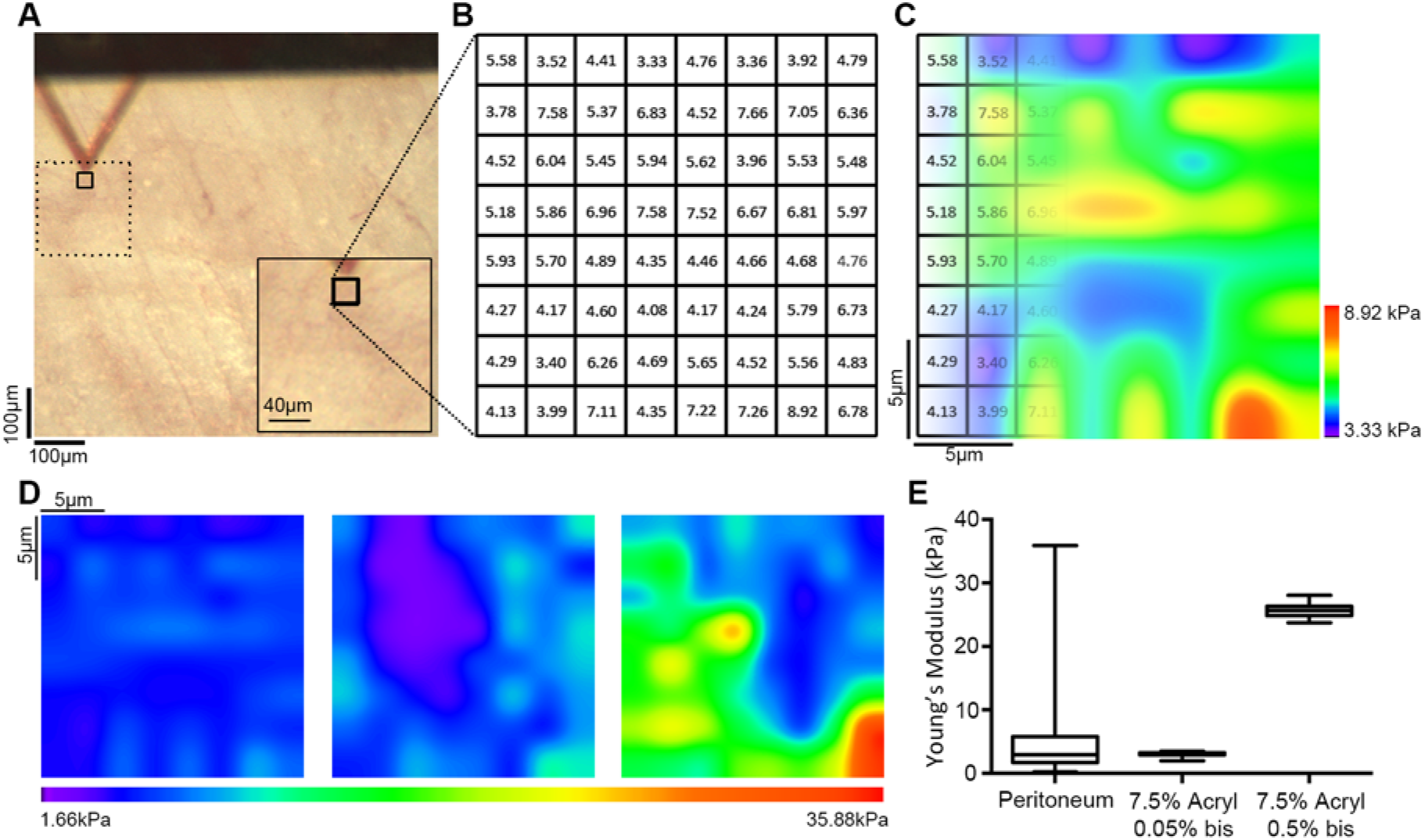
Assessment of peritoneal rigidity by atomic force microscopy. (A) Mouse peritoneum was mounted on an atomic force microscope (AFM) and force-indentation curves were obtained using a 5 μm spherical probe mounted on a cantilever (triangle). (B) Force-maps were generated by obtaining an 8 x 8 grid of Young’s elastic modulus (E) values from 20 μm x 20 μm regions within a tissue section. (C and D) The grids were transformed into color-coded heat maps, the scales of which could be was set using minimum and maximum values from an individual map (C) or from values from a series of maps to compare results between numerous regions across several different animals (D; note that the leftmost map in (D) is the same map, shown with different value scaling, as in (C)). (E) Box and whisker plots showing minimum and maximum (lower and upper whiskers), 25th-75th quartiles (box), and median (horizontal line inside box) E values from normal mouse peritoneum (ms. peritoneum) and soft or stiff polyacrylamide hydrogels (0.05% or 0.5% bis-acrylamide crosslinker, n = 1724 for mouse peritoneum and n = 128 for each gel).

### Cell size, F-actin organization, and focal adhesion formation positively correlate with substrate rigidity in SKOV3 human EOC cells

While studies have shown that cell spreading and morphology as well as focal adhesion maturation are positively correlated with the rigidity of the ECM in a variety of cells^31–39^, this correlation has not been investigated in EOC cells. To this end, SKOV3 cells – derived from a human ovarian adenocarcinoma – were plated on fibronectin-coated ‘soft’ (3 kPa) or ‘stiff’ (25 kPa) gels and on fibronectin-coated glass coverslips (Young’s modulus = ~70 GPa). Fibronectin was chosen as an adhesive substrate owing to its abundance in ovarian tumor-associated stroma^40^ and its well-documented importance in the migration, invasion, and metastasis of ovarian cancer^40–48^. After overnight incubation, the cells were fixed and stained to visualize F-actin, and paxillin (Figure 2A). Striking morphological changes were readily observed in cells plated on substrates of varying rigidity. Specifically, cells plated on glass, stiff 25 kPa gels and soft 3 kPa gels spread with significantly different average areas of 3920 μm^2^, 1673 μm^2^, and 111 μm^2^, respectively (Figure 2B). Although cells plated on soft gels did not spread significantly, they were stably adherent, as changing the media and processing for immunofluorescence did not dislodge them. Also, while no higher-order actin cytoskeletal structures were apparent in cells on soft gels, cells plated on stiff gels had clear stress fibers, peripheral actin cables, and leading edge ruffles, while cells on glass had numerous, robust stress fibers (Figure 2A). The number, size, and aspect ratio of paxillin-containing focal adhesions also increased with increasing substrate stiffness (Figure 2C-2E). These data demonstrate that EOC cell size, actin cytoskeletal organization, and focal adhesion morphology positively correlate with matrix stiffness.

**Figure 2.**
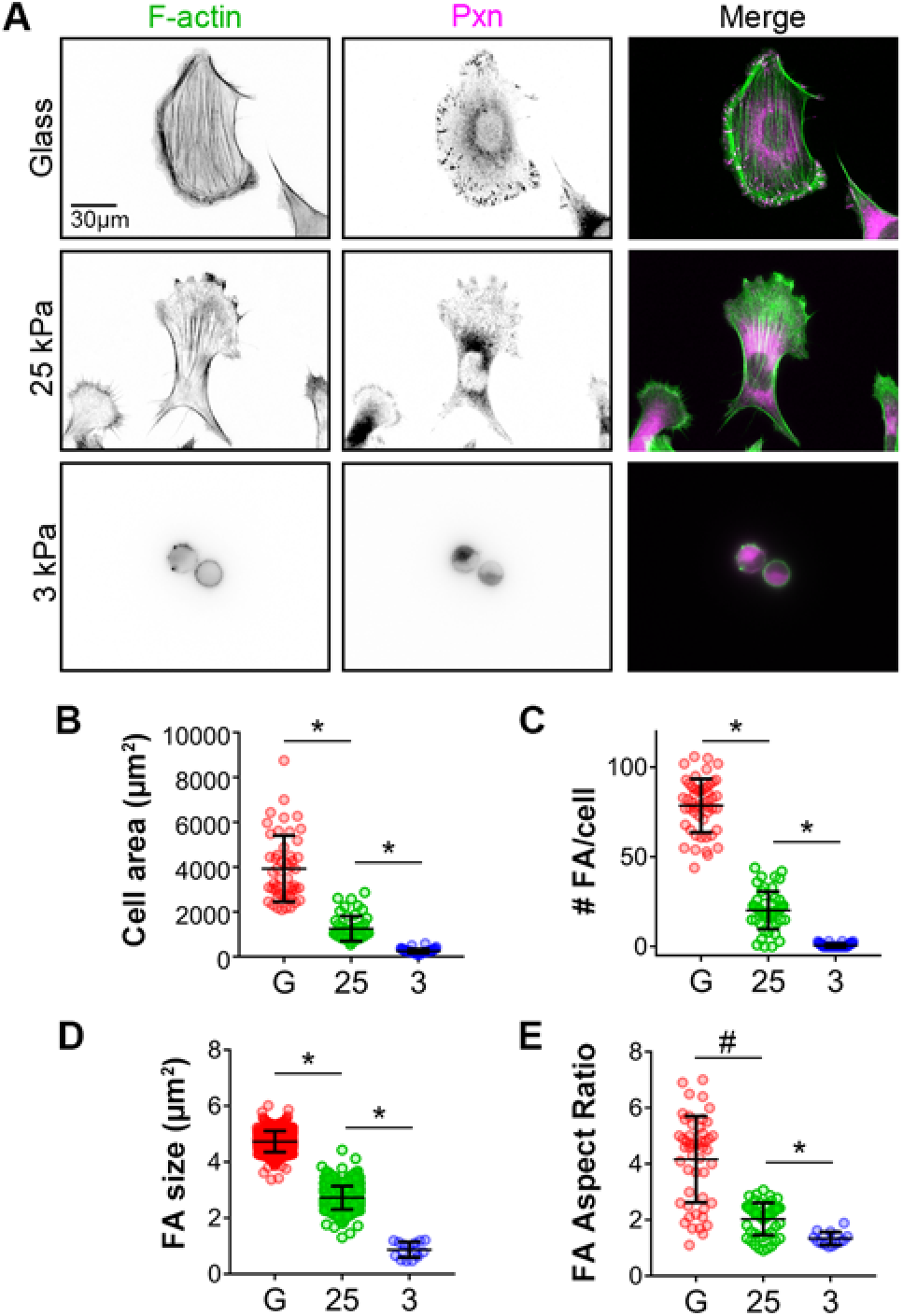
Cell size and focal adhesion morphology positively correlate with substrate rigidity. (A) SKOV-3 cells plated on fibronectin coated glass (top), 25 kPa, and 3 kPa hydrogels (middle and bottom) were fixed and stained with phalloidin and paxillin (Pxn) antibody to visualize F-actin stress fibers and focal adhesion complexes (FA) respectively. (B-E) Immunofluorescence images were analyzed for cells on each substrate to determine cell size (B), number of FA/cells (C), FA size (D), and FA aspect ratio (E). Graphs show all measured values as colored symbols as well as mean values ± s.d. as black bars (n = 50 cells (B and C) and 750 or 15 FA for glass and 25 kPa or 3 kPa, respectively (C and D); *, *p* <0.0001 and #, *p* = 0.0261, using the Mann-Whitney test).

### Myosin Light Chain phosphorylation in SKOV3 EOC cells is regulated by ECM tension

There is a well-established reciprocity between focal adhesion formation and actomyosin contractility in the context of mechanotransduction. The stiffness-dependent increase in focal adhesion number, size, and aspect ratio suggested that actomyosin contractility in EOC cells might also be increased by substrate rigidity. Actomyosin contractility regulates cell shape and the dynamics of the actin cytoskeleton and focal adhesions and is also required for many of the cellular responses to mechanical cues^1,2,49,50^. Phosphorylation of the myosin regulatory light chain (MLC) and assembly of myosin mini-filaments onto F-actin are important facets of generating cellular contractility and regulating tensional homeostasis^51,52^. To begin to assess the effect of substrate rigidity on EOC contractility, SKOV3 cells adherent to FN-coated glass, 25 kPa, and 3 kPa gels were also stained to visualize active, phosphorylated MLC (containing phospho-Thr18 and phospho-Ser19). Visualization and quantification of phospho-MLC staining revealed that the majority of active, phosphorylated MLC co-localized with F-actin in cells on 25 kPa gels and on glass (Figure 3A). In contrast, there was significantly less colocalization between phospho-MLC and F-actin in cells plated on 3 kPa gels (Figure 3A). Furthermore, immunoblot analysis of lysates generated from cells plated either on glass or hydrogels of varying rigidity showed a positive correlation between MLC phosphorylation and substrate rigidity (Figure 3B & C). Together, these data show that, in EOC cells, myosin activity increases with increasing matrix stiffness.

**Figure 3.**
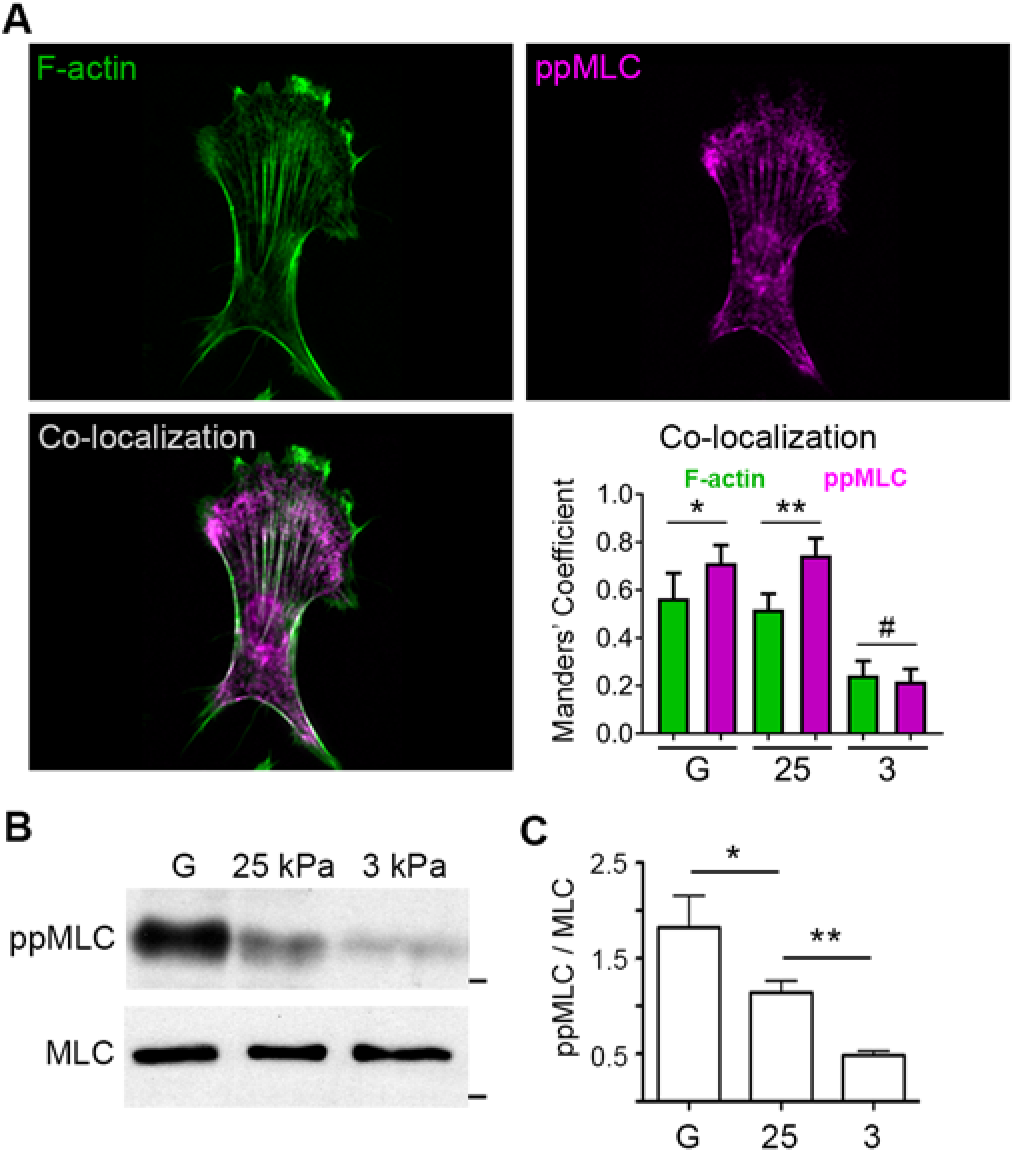
Substrate rigidity enhances actin-myosin co-localization and myosin phosphorylation. (A) SKOV-3 cells plated on glass (G), 25 kPa, or 3 kPa hydrogels were fixed and stained to visualize F-actin and dually-phosphorylated myosin light chain (ppMLC). Images of a representative cell on 25 kPa hydrogel are shown (*n.b*. this is the same cell shown in Figure 2). Co-localization of F-actin and ppMLC was assessed by pixel-by-pixel image correlation analysis performed using intensity correlation analysis to produce Mander’s correlation coefficients. Average Mander’s coefficients are plotted as mean ± SEM (n= 15 cells; *, *p* = 0.0015 using Wilcoxon test; **, *p* < 0.0001 using Wilcoxon test; #, no significant difference between Manders’ coefficients for F-actin and ppMLC on 3 kPa gels, while the coefficients for both F-actin and ppMLC on 3 kPa are significantly different (*p* < 0.0001, unpaired t-test) than the corresponding values on 25 kPa and glass). (B) Lysates from SKOV-3 cells plated on the indicated substrates were separated by SDS-PAGE and probed with antibodies against ppMLC (Thr18/Ser19) and total myosin light chain (MLC). (C) The relative ratio of ppMLC to total MLC was determined by densitometry (mean ± s.d.; n=3; *, *p* < 0.05; **, *p* < 0.01).

### Cellular traction forces positively correlate with substrate rigidity in SKOV3 cells

The positive correlation between myosin activity and substrate stiffness suggested that cellular contractility and tension might follow the same relationship. Thus, we determined the effect of matrix rigidity on traction forces in SKOV3 cells. Given that glass surfaces are not amenable to traction force microscopy, we employed rigid 125 kPa in addition to the stiff 25 kPa and soft 3 kPa gels for these and subsequent experiments, in order to extend the experimental correlation range. As predicted from the aforementioned myosin data, cellular traction (mean, maximum, and total) was low in cells on pliable 3 kPa gels and increased significantly in cells on 25 kPa and 125 kPa gels (Figure 4A-D), indicating that EOC cells exhibit tensional homeostasis – increasing their contractility and counter-tension as the stiffness of their underlying matrix increases.

**Figure 4.**
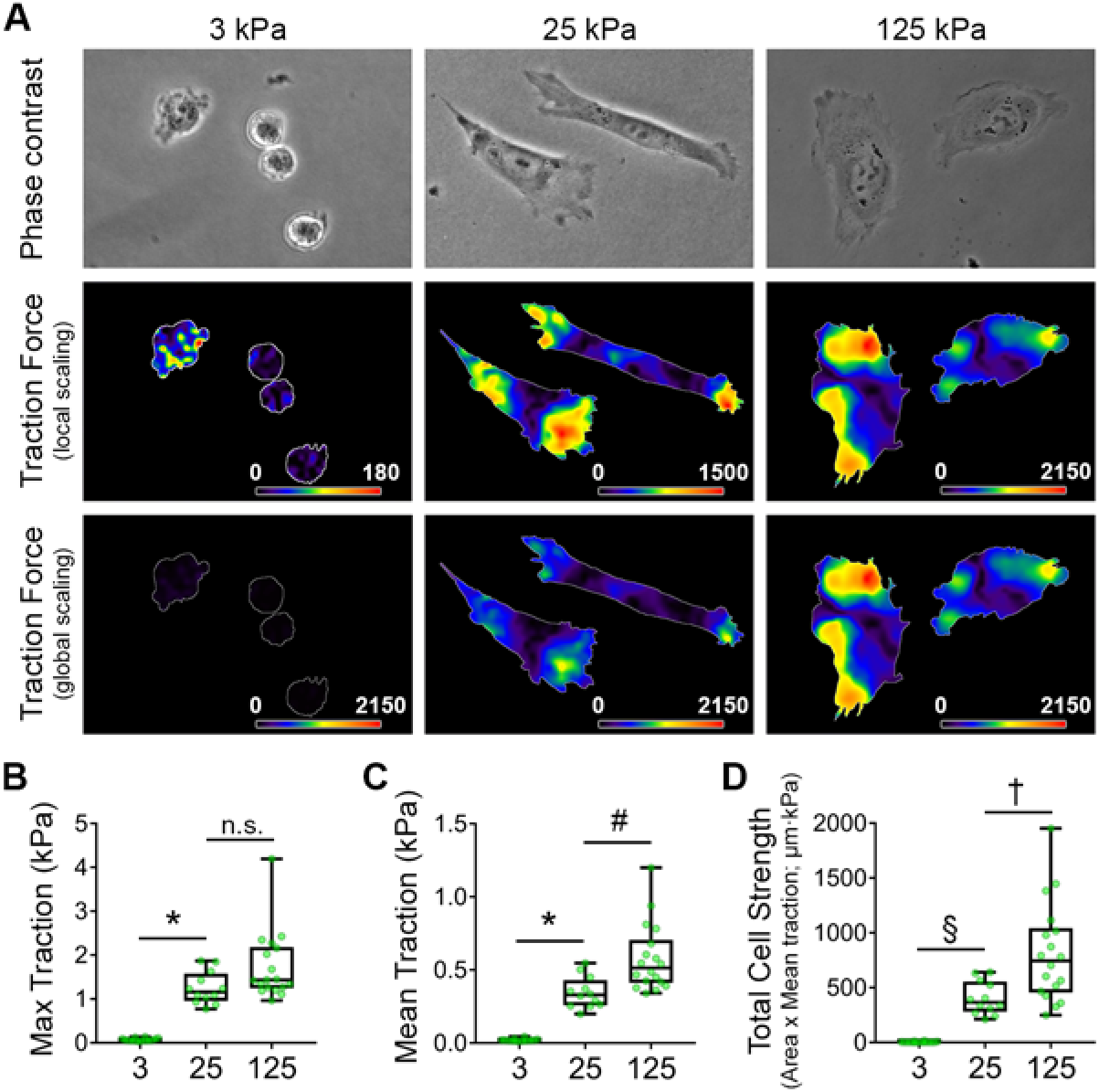
Cellular traction forces positively correlate with substrate rigidity. (A) Traction force maps for SKOV-3 cells plated on fluorescent nanosphere-functionalized hydrogels were generated using Particle Image Velocimetry (PIV) and Fourier-Transform Traction Cytometry (FTTC) plugins for ImageJ. Phase contrast images (top) and traction force maps (middle and bottom) shown for representative cells adhered to gels of the indicated of the indicated Young’s modulus values. Traction force maps are shown with both local (middle) and global (bottom) scaling to highlight differences in traction forces across substrate rigidity. (B-D) Maximum traction (B), mean traction (C), and total cell strength were measured for cells adhering to hydrogels of each elastic modulus. Graphs depict all of the data points (green symbols), with box & whisker plots showing maximum & minimum values (whiskers), the 25^th^ to 75^th^ percentiles (box), and the median value (box line). (n = 15, 12, and 18 cells from 3 experiments for 3, 25, and 125 kPa samples, respectively; *, *p* < 0.0001; #, *p* = 0.007; §, *p* = 0.0037; †, *p* = 0.002; n.s. = not significant).

### Matrix rigidity enhances SKOV3 cell migration

In addition to cell morphology and traction force, cell migration has also been shown to be enhanced by matrix rigidity^53–58^. Thus, to examine this relationship in the context of EOC, we seeded SKOV3 cells on soft, stiff, and rigid hydrogels (3, 25, and 125 kPa, respectively) and tracked their random migration over a period of 14 h. Less than 1% of cells plated on soft (3 kPa) gels showed significant migration, defined as a maximum displacement > 50 μm (Figure 5A & B), and the few cells that did migrate barely exceeded this threshold (Figure 5C and Supplemental Movie S1, top panel). In contrast, 40-50% of cells plated on stiff (25 kPa) or rigid (125 kPa) substrates showed significant migration, with mean displacements of ~120 μm and maximum displacements in excess of 300 μm (Figure 5A-C and Supplemental Movie S1, middle and bottom panels). These data show that the rigidity-dependent increases in EOC cell area, cytoskeleton & adhesion organization, and traction force collaborate to support more efficient cell migration.

**Figure 5.**
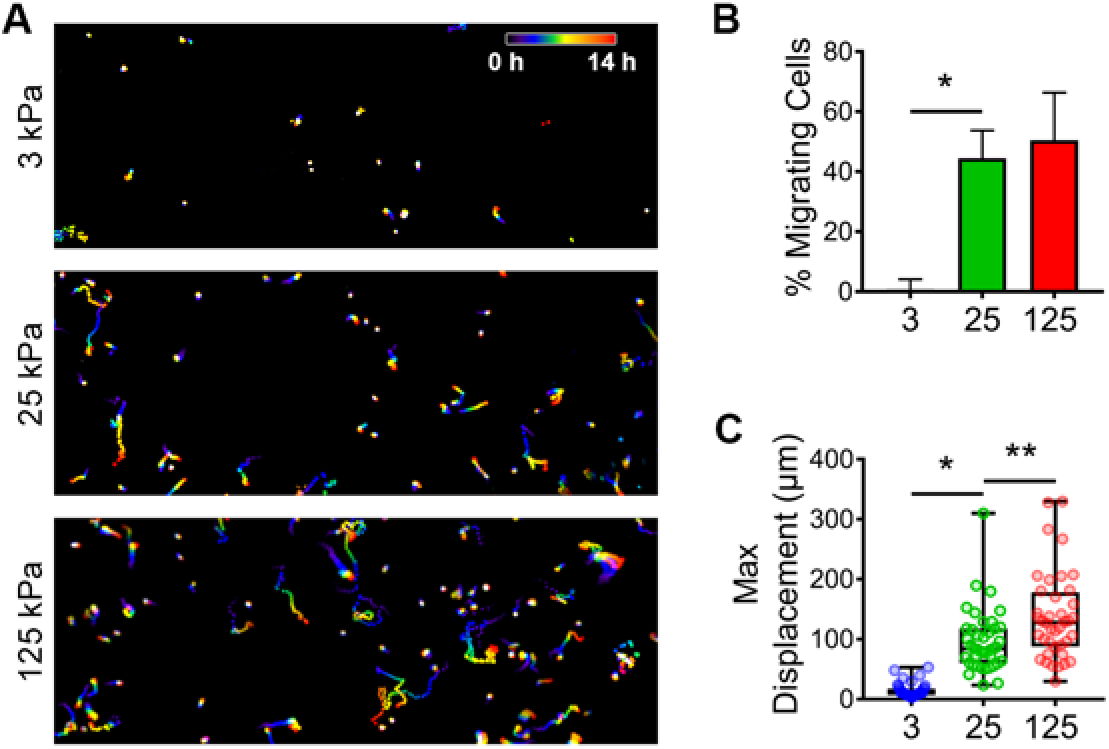
Substrate rigidity regulates ovarian cancer cell migration. (A) SKOV-3 cells plated on 3, 25, and 125 kPa hydrogels were monitored by live-cell microscopy (capture rate = 4 frames / h) and nuclei were tracked using AIVIA software. Consecutive frames of time-lapse movies of were color-coded (as depicted by the “0 h – 14 h” scale bar) and maximum intensity projections of the stacks were generated to show the cumulative cell tracks over 14 h. (B) Maximum displacements were calculated by determining the largest Euclidean distance between the position of each cell at t = 0 and its position at each frame of the time lapse image. The graph depicts all of the data points (colored symbols), with box & whisker plots showing maximum & minimum values (whiskers), the 25^th^ to 75^th^ percentiles (box), and the median value (box line). (n = 40 cells from 3 experiments; *, *p* < 0.0001; **, *p* = 0.002 using a Mann-Whitney test). (C) The percent of migrating cells was calculated by determining the number of cells with a maximum displacement > 50 μm versus the total number of cells per movie. (n = 79, 159, and 180 cells from 3 experiments for 3, 25, and 125 kPa samples, respectively; *, *p* = 0.008).

### EOC cells exhibit durotaxis

We postulated that if EOC cell migration is modulated by static, bulk differences in substrate stiffness, as described above, then acute, localized and/or directional changes in matrix rigidity might also influence EOC cell migration. Indeed, spatially discrete changes in matrix stiffness have been shown to induce migration in the direction of increased stiffness numerous cell types^53,59^. This process, termed durotaxis, has been demonstrated to be a crucial regulator of cell migration and its myriad dependent processes^49,60,61^.Though durotaxis has been observed and characterized in other cell types, it has yet to be described in the context of EOC. To this end, randomly migrating SKOV3 cells plated on stiff (25 kPa) hydrogels were exposed to a durotactic stimulus by deforming the hydrogel near a given target cell with a glass pipet as described in *Methods* (Figure 6A) and phase microscopy images were acquired before stretch and for 80 min after stretch. SKOV3 cells exhibited a rapid and robust durotactic response when subjected to mechanical stretch (Figure 6B and Supplemental Movie S2). Quantification of the turn angle in response to directional stretch, as compared to the angles formed by unstretched cells over the same time period, confirmed that the high degree of directionality of stretch-induced migration was not stochastic but rather driven by the directional increase in matrix tension (Figure 6C and D). These results show, for the first time, that EOC cells exhibit a durotactic response to directional increases in matrix tension.

**Figure 6.**
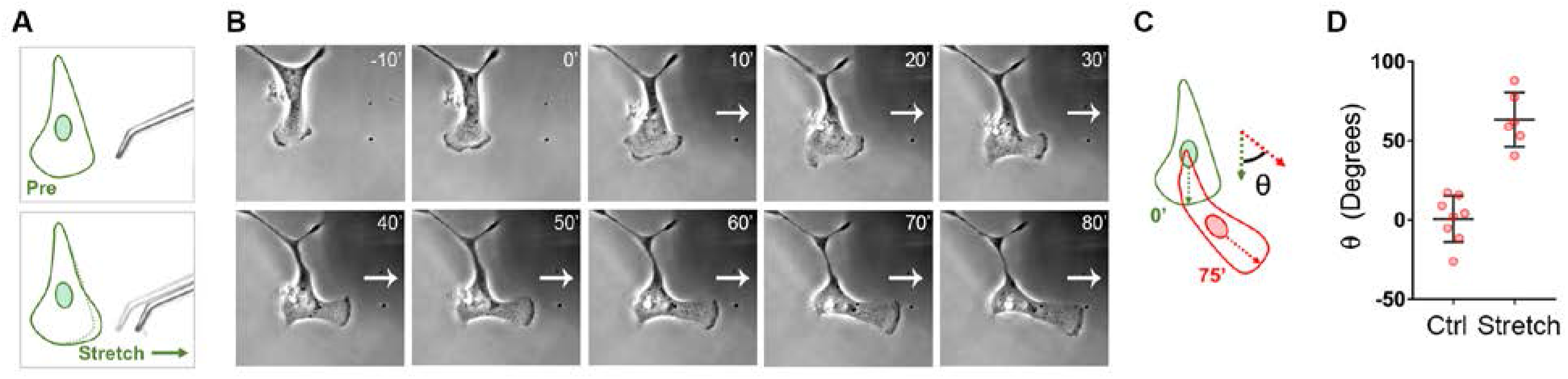
Ovarian cancer cells exhibit durotaxis. (A) SKOV3 cells plated on hydrogels with rigidity of 25 kPa were allowed to adhere overnight before hydrogels were deformed with a glass micropipette and pulled orthogonal to a single migrating cell’s established direction of travel. Stretch was maintained over a period of 80 minutes with images taken every minute. (B) Example of a positive durotactic response. Time lapse imaging shows morphology and position of the SKOV3 cell at 10 minutes prior to stretch, immediately before stretch, and every ten minutes during stretch for 80 minutes. White arrow indicates direction of maintained stretch. (C) Durotactic response quantified as the turn angle (θ) determined as the angle of deflection from the original direction of travel after 75 minutes. Direction of travel at each time point established by a line drawn from the nucleus to the leading edge. (D) Quantification of turn angle of responding SKOV3 cells presented with durotactic stretch for 75 minutes compared to a similar number of control (Ctrl) cells not presented with a stimulus. Graph depicts all data points (red symbols) as well as the mean turn angle (± s.d.; n = 8 and 5 cells for control and stretched cells, respectively; *p* = 0.007 by the Mann-Whitney test).

### Matrix rigidity enhances the disaggregation of multicellular EOC spheroids

We reasoned that if the migration of individual EOC cells is influenced by the mechanical properties of their microenvironment, then the motile dynamics of multicellular EOC spheroids might also be regulated by changes in ECM tension. Studies have shown that spheroids are the minimal metastatic unit of EOC^20,24^, with spheroid formation, adhesion, and/or disaggregation being critical for colonization of the peritoneum and commonly used as readouts of EOC metastatic potential^24,25,27,62^.To date, however, no studies have shown the effects of matrix stiffness on spheroid adhesion or disaggregation.

To investigate this, multicellular spheroids were formed by culturing SKOV3 cells in agarose-coated microwells as described in *Methods*, then re-plated onto FN-coated hydrogels of varying rigidities and allowed to attach and disaggregate overnight. Spheroids plated onto soft 3 kPa gels attached loosely within 2 h but showed no disaggregation by 20 h (Figure 7A and B). In contrast, spheroids on stiff 25 kPa gels adhered more tightly and started to lose their sphericity by 2 h after plating and showed significant disaggregation (with an average 6.6-fold increase in their ‘footprint’ or spread area) at 20 h (Figure 7A and B). Spheroids plated on rigid 125 kPa gels adhered and started to disaggregate by 2h and showed an average 8.5-fold increase in spread area by 20 h (Figure 7A and B). These results demonstrate, for first time, that the mechanical properties of the underlying substrate directly regulate EOC spheroid disaggregation and in turn suggest that stiffening of ECM in the tumor microenvironment may play a role in EOC implantation and invasion *in vivo*.

**Figure 7.**
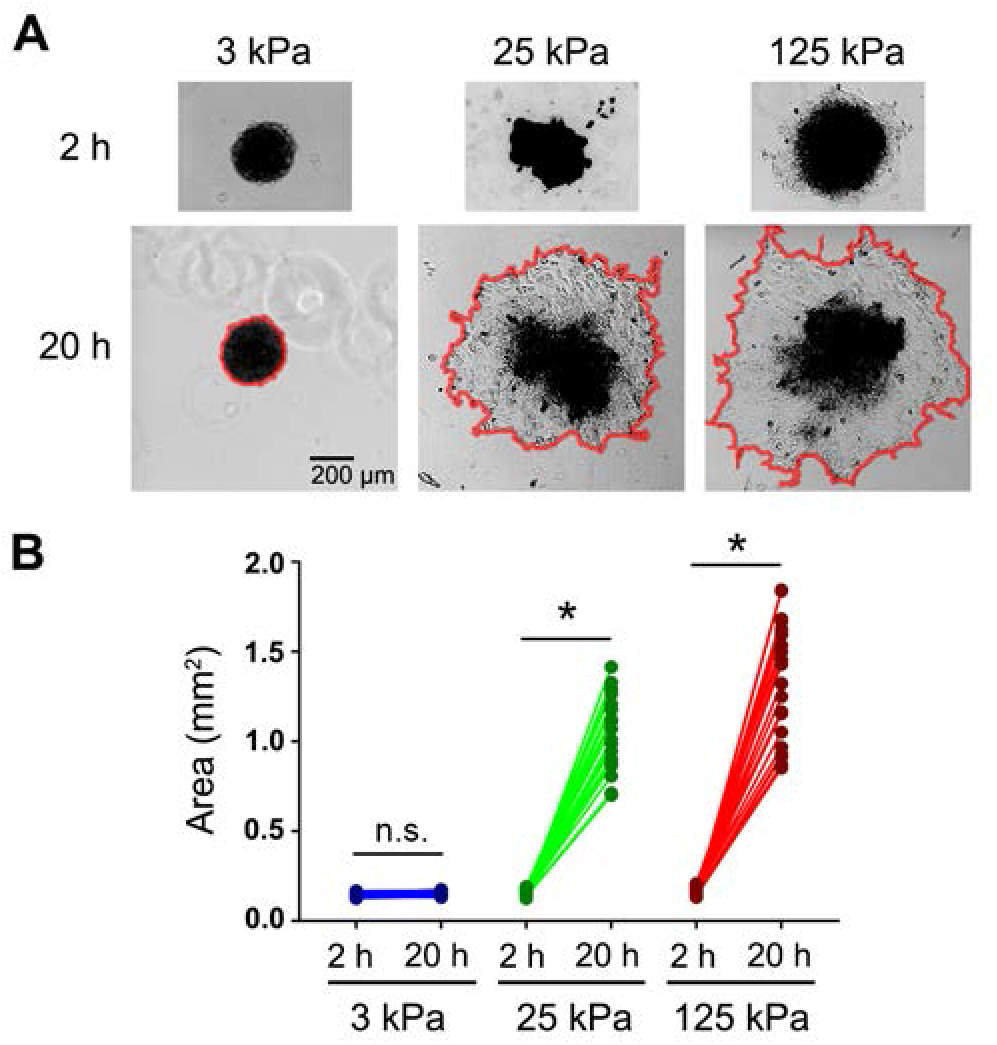
Disaggregation of ovarian cancer spheroids is dependent on substrate rigidity. (A) SKOV3 multicellular spheroids were plated on hydrogels with rigidities of 3 kPa, 25 kPa, and 125 kPa and imaged at 2 h and 20 h post plating. Red lines in the 20 h panels help visualize the boundaries of the tumor spheroids. (B) The graph depicts the change in area of each spheroid measured at 2 h and 20 h by manually outlining spheroids at the indicated time points and calculating the area using ImageJ (n = 24 spheroids for each condition; *, *p* < 0.0001; n.s. = not significant).

### A mechanotransduction pathway comprising ROCK, actomyosin contractility, and FAK controls disaggregation of ovarian cancer spheroids

The aforementioned effect of matrix rigidity on EOC spheroid disaggregation indicates that disaggregation is dependent on and driven by mechano-sensitive biochemical pathways and suggests that disruption of such pathways might inhibit disaggregation. To begin to investigate this, we first examined the effect of inhibiting the activity of canonical biochemical components of mechanotransduction at the single-cell level by assessing cellular traction force, focal adhesion morphology and focal adhesion kinase (FAK) phosphorylation, and the morphology and intensity of F-actin and phospho-MLC staining in cells on stiff hydrogels.

As discussed above for Figure 3, myosin is a principal component of cellular contractility. As expected, then, inhibition of the ATPase activity of non-muscle myosin II with blebbistatin all but eliminated cellular traction force (Figure 8A and D) and altered focal adhesion morphology (Figure 8E), without demonstrable effect on FAK phosphorylation at Tyr397 (Figure 8E and F). Furthermore, while blebbistatin treatment dissolved F-actin stress fibers (Figure 8G), it did not affect the intensity of phospho-MLC staining or its co-localization with residual F-actin structures (Figure 8G and H).

**Figure 8.**
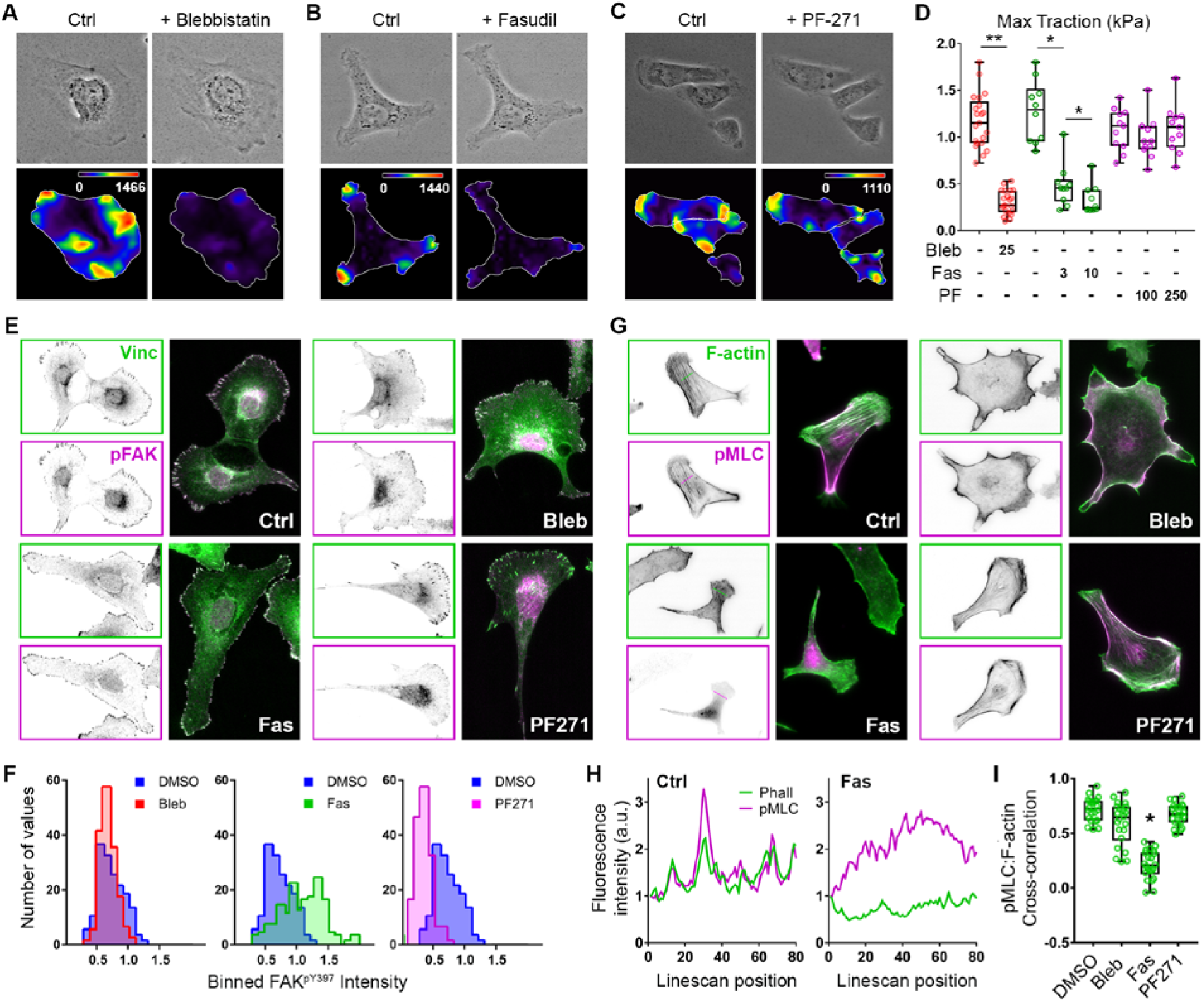
Cellular traction forces are dependent on ROCK activity and actomyosin contractility but independent of FAK activity. (A-C) Phase contrast images (top) and traction force maps (bottom) calculated for SKOV3 cells adhered to 25 kPa hydrogels following 30 minute treatment with 25 μM blebbistatin (A), 10 μM fasudil (B), and 0.25 μM PF-271 (C). (D) Maximum traction forces were analyzed following 30 minute treatment with 0.1% DMSO or the indicated compound (μM) (n = 10, 11, or 21 cells for blebbistatin, fasudil, or PF-271, respectively; *, p = 0.002; **, p < 0.0001 using the Wilcoxon matched pairs test). (E) SKOV3 cells plated on 25 kPa hydrogels were treated with 0.1% DMSO (Ctrl), 25 μM blebbistatin (Bleb), 10 μM fasudil (Fas), or 0.25 μM PF-562271 (PF271) for 30 minutes before being fixed and stained to visualize vinculin (Vinc) and FAK phosphorylated on Tyr397 (pFAK; pY397). (F) Individual focal adhesions were outlined by thresholding vinculin images and running a custom macro in ImageJ which transferred each focal adhesion ROI to the raw vinculin and pFAK images and measured each signal intensity. Mean pFAK intensity was normalized to mean Vinc intensity within each adhesion. Histograms represent binned, normalized pFAK levels within adhesion for each treatment compared to DMSO control (n > 800 adhesions from > 8 cells per treatment). (G) SKOV3 cells were plated and treated as in panel (E) then stained for F-actin and phospho-MLC. (H) Linescans of relative F-actin and phospho-MLC intensity (in arbitrary units (a.u.)) for representative control and fasudil treated cells were plotted from the dotted lines shown in (G). (I) Pearson’s cross-correlation analysis of phospho-MLC and F-actin intensity linescans as shown in (H). The graph depicts all of the data points (green symbols), with box & whisker plots showing maximum & minimum values (whiskers), the 25^th^ to 75^th^ percentiles (box), and the median value (box line) (n = 22 linescans (1 scan/cell) from 3 experiments; *p* < 0.0001 using the Kruskal-Wallis test with multi-comparisons (unannotated differences are not significant)).

Also as discussed earlier, actomyosin contractility is regulated by phosphorylation of the MLC through a complex network of kinases and phosphatases. An important component of this network are the ROCKs (Rho-dependent coiled-coil-containing kinase, or Rho-kinase), which couple activation of the Rho GTPase to MLC phosphorylation through direct (*via* phosphorylation of MLC) and indirect (*via* phosphorylation and inhibition of MLC phosphatase) pathways to promote cell-matrix tension^52,63^. Like blebbistatin, treatment of SKOV3 cells with fasudil (also known as HA-1077), a ROCK inhibitor that has previously been shown to inhibit EOC cell invasiveness *in vitro* and to reduce intraperitoneal tumor burden and ascites formation *in vivo*^64^, also led to a substantial reduction in traction force (Figure 8B and D). Focal adhesion size and aspect ratio were also reduced by fasudil treatment, although with a concomitant and paradoxical increase in phospho-FAK levels (Figure 8E and F); the reason for this increase is not known. Finally, unlike blebbistatin but as expected, inhibition of ROCK with fasudil essentially eliminated cytosolic phospho-MLC staining and coincidence with F-actin (Figure 8G-I).

A central mediator of adhesion-dependent signaling, the role of FAK in mechanotransduction is regarded as being less in the generation of cellular tension and more in the transduction of signals downstream of mechanical cell-matrix interactions^65–67^. Nonetheless, its demonstrated importance in both functional mechanical signaling^68–71^ and in the biology of EOC^72–77^ prompted us to assess the effect of FAK inhibition on the EOC mechano-response. Unlike blebbistatin and fasudil, treatment of SKOV3 cells with PF-271 (also known as PF-562271), a potent and highly selective inhibitor of FAK, had no significant effect on cellular traction force (Figure 8C and D) or stress fiber and phospho-MLC architecture (Figure 8G and I), but significantly reduced phospho-FAK intensity while preserving or slightly enhancing focal adhesion size and aspect ratio (Figure 8E and F), consistent with FAK’s role in focal adhesion turnover^78^.

To further assess the mechanical regulation of spheroid disaggregation, we first examined the traction forces along the periphery of spheroids adhered for 2 h to soft, stiff, or rigid FN-coated hydrogels. Consistent with their respective degrees of disaggregation, spheroids on 3 kPa gels exerted low traction forces on their substrate, while spheroids on 25 kPa and 125 kPa gels exhibited substantial traction forces, especially those exerted by the vanguard cells at the periphery of the disaggregation front (Figure 9A). We then examined the effect of inhibiting the aforementioned regulators of mechanotransduction on EOC spheroid disaggregation on stiff 25 kPa hydrogels. Despite intersecting the mechanotransduction machinery at different nodes (as detailed in Figure 8), inhibition of myosin, ROCK, and FAK (*via* blebbistatin, fasudil, and PF271, respectively) all dramatically inhibited SKOV3 spheroid disaggregation (Figure 9B and C). These data, combined with those presented in the previous figure, indicate that FAK functions downstream of actomyosin contractility and cellular tension in a mechanotransduction pathway that regulates EOC spheroid disaggregation.

**Figure 9.**
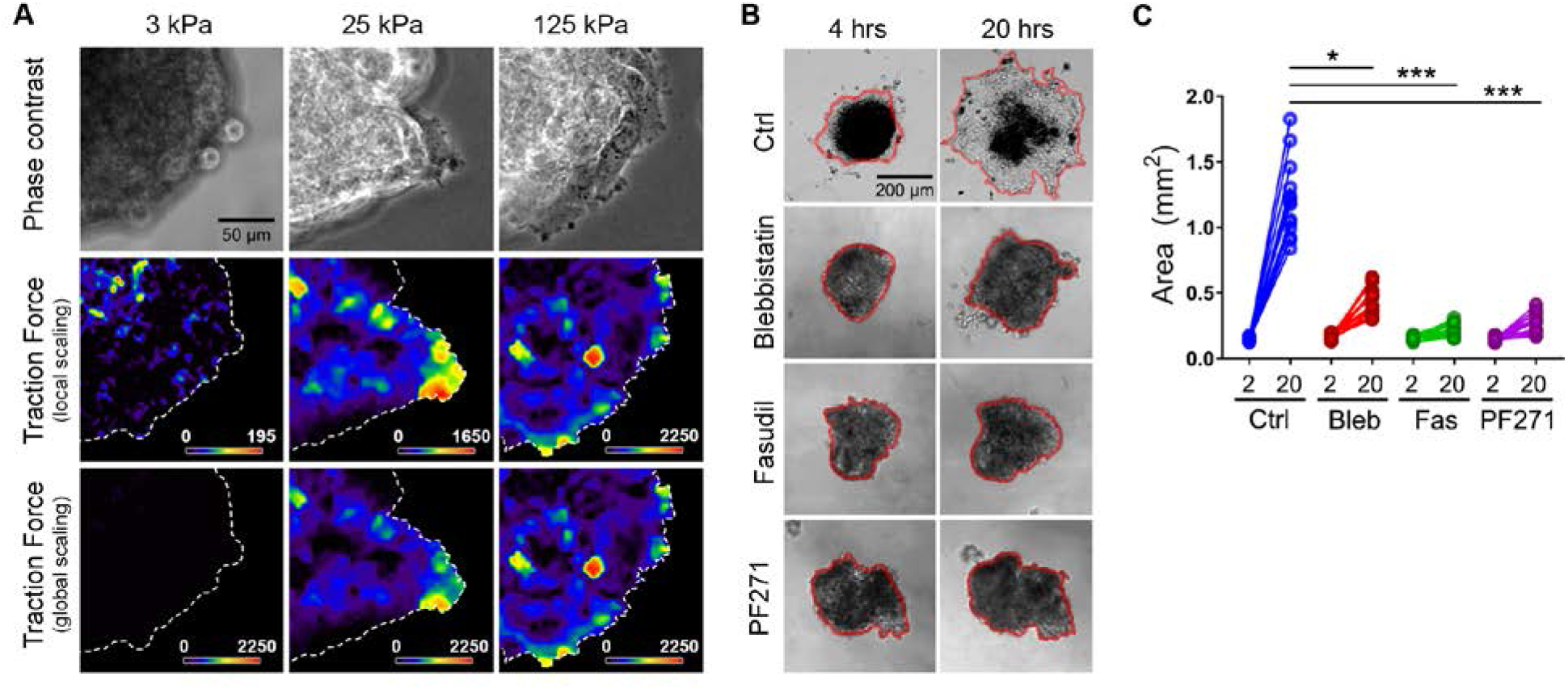
A mechanotransduction pathway comprising ROCK, actomyosin contractility, and FAK controls disaggregation of ovarian cancer spheroids. (A) Traction forces underlying SKOV3 ovarian cancer spheroids at 2 h after plating on hydrogels with rigidities of 3 kPa, 25 kPa, or 125 kPa, as indicated. Phase contrast images (top), calculated traction forces with local scaling (middle), and calculated traction forces with global scaling (bottom) are shown for representative spheroids on each hydrogel rigidity. (B) Phase contrast images of SKOV3 ovarian cancer spheroids, treated 2 h after plating with 0.1% DMSO (Ctrl), 25 μM blebbistatin, 10 μM fasudil, and 0.25 μM PF271, were captured at 4 h and 20 h post plating on hydrogels with rigidity of 25 kPa. Red lines help visualize the boundaries of the tumor spheroids at each timepoint. (C) Quantification of changes in area between 2 hours and 20 hours of DMSO control-, blebbistatin-, fasudil-, and PF271-treated spheroids (n = 16 spheroids for each treatment; *, *p*= 0.0286; ***, *p* < 0.001).

### Substrate rigidity controls nuclear localization of YAP1 in SKOV3 cells

Given the present data establishing a positive correlation between matrix stiffness and cellular behaviors related to EOC invasion and metastasis, we endeavored to investigate whether matrix stiffness regulated other characteristics associated with aggressive disease. YAP1 is a transcriptional co-activator downstream of the Hippo signaling pathway that is important both in normal development (e.g. for controlling cell stemness and organ size and development) and in initiation, progression, and metastasis of many cancers^79,80^. YAP1 is also widely regarded as a mechanotransducer, as its transcriptional activity is responsive to a complex array of mechanical and geometric cues including the shape, density, and polarity of cells as well as the architecture of the actin cytoskeleton and the mechanics of the microenvironment^79,81,82^. For example, YAP1 nuclear localization is regulated by substrate rigidity, although the mechanisms connecting cell-matrix tension and YAP1 nucleocytoplasmic shutting are not fully understood^82–84^. Importantly, *YAP1* is a *bona fide* ovarian cancer oncogene, and the expression, activity, and nuclear localization of the YAP1 gene product is associated with aggressive disease and poor prognosis^85–88^. Thus, we investigated whether YAP1 nuclear localization in EOC cells was affected by matrix rigidity. SKOV3 cells were plated onto FN-coated hydrogels of increasing rigidities, as well as onto FN-coated glass coverslips (as a ‘positive control’ for maximum substrate stiffness) overnight, then fixed and stained to visualize the actin cytoskeleton, the nucleus, and YAP1 localization. As expected, cells plated on glass were well-spread and polarized, contained prominent actin stress fibers, and showed robust nuclear localization of YAP1 (Figure 10A-C), although this localization was dependent on actin cytoskeletal integrity rather than actomyosin contractility (Supplemental Figure S1), in agreement with recent observations in other cells^84^. In contrast to cells on glass, the rounded cells on soft 3 kPa gels showed peak YAP1 staining that was effectively excluded from the nucleus in (Figure 10A and B) and had a low overall nuclear: cytoplasmic ratio of YAP1 localization (Figure 10C). Importantly, nuclear localization of YAP1 and the overall nuclear:cytoplasmic ratio of YAP1 intensity increased proportionally in cells plated on 25 kPa and 125 kPa gels (Figure 10A-C). These data demonstrate that, in addition to motile behavior, substrate rigidity also positively regulates the nucleocytoplasmic shuttling of YAP1, a mechano-responsive enhancer of EOC aggression and metastasis.

**Figure 10.**
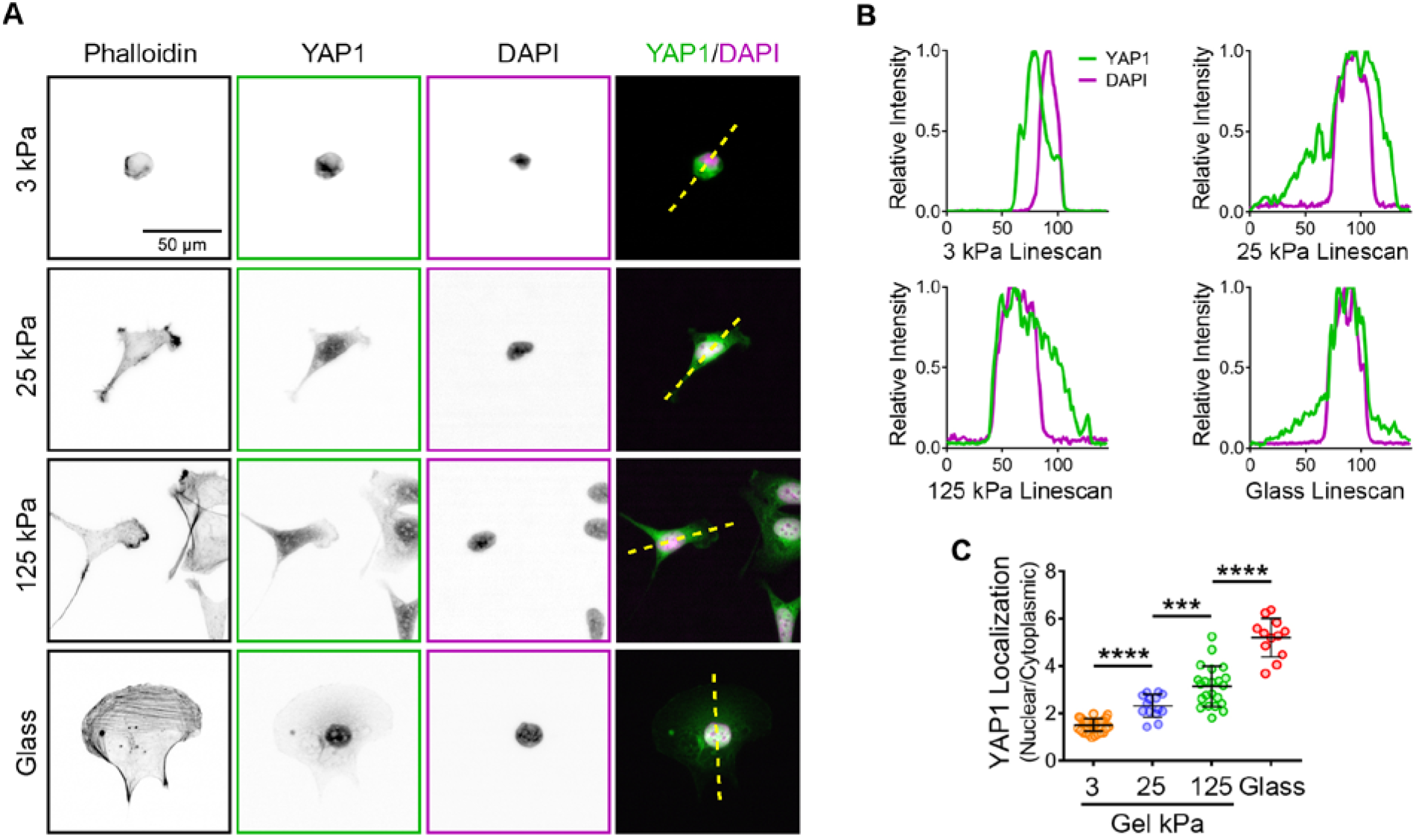
Substrate rigidity controls nuclear localization of YAP1 in ovarian cancer cells. (A) SKOV3 cells plated on hydrogels with elastic moduli of 3 kPa, 25 kPa, and 125 kPa, or on glass coverslips, as indicated, were allowed to adhere for 18 h before being fixed and stained with phalloidin, YAP1 antibody, and DAPI to visualize F-actin, YAP1, and nuclei, respectively. Yellow dotted lines through the cells in the overlay images (last column) were used to generate the intensity linescans shown in the next panel. (B) Plotted intensity of YAP1 and DAPI signals through the linear regions of interest depicted in (A). The y-axes represent relative fluorescence intensity of the indicated signals and the x-axes represent the position along the linear ROI. (C) Quantification of YAP1 localization given as a ratio of YAP1 signal in the nucleus to that in the cytoplasm for cells on the indicated substrates. The graph depicts all measure values (colored symbols) and the mean values (± s.d.; n = 34, 14, 22, and 16 cells for 3 kPa, 25 kPa, 125 kPa, and glass, respectively; ***, *p* = 0.0025; ****, *p* < 0.0001 using Mann-Whitney tests).

### Mechanical regulation of cell morphology, migration, tension, and spheroid disaggregation is maintained in highly metastatic SKOV3.ip1 cells

Collectively, the present work indicates that increasing substrate rigidity enhances characteristics of EOC disease progression and spread (i.e. migration, spheroid disaggregation, YAP1 nuclear localization). However, a previous report has suggested that metastatic potential in EOC cells correlates with preferential cell spreading and migration on soft matrices^89^. To examine whether the positive correlation between matrix stiffness and metastasis-related cell behaviors was reversed or altered with increasing metastatic potential, we assessed hallmark aspects of mechano-responsiveness in SKOV3.ip1 cells, a subclone of SKOV3 cells derived by serial selection and expansion of intraperitoneal metastases^72^. As seen for SKOV3 cells, there was for highly metastatic SKOV3.ip1 a positive correlation between matrix stiffness and cell spreading (Figure 11A), traction forces (Figure 11B-D), and random cell migration (Figure 11E-G). Importantly, SKOV3.ip1 cells also exhibited a robust durotactic response (Figure 11H and I; Supplemental Movie S3), which dramatically and unequivocally demonstrates the positive correlation between matrix stiffness and migration in these cells. Finally, we assessed the influence of substrate stiffness on disaggregation of SKOV3.ip1 spheroids. As noted in *Methods*, this cell line did not exhibit the same kinetics of spheroid formation as the parental SKOV3 line, requiring 5-7 d to tightly aggregate into numerous, small spheroids and 14-18 days to form larger, cohesive aggregates (as opposed to the essentially complete compaction into a single spheroid within 3 d seen for SKOV3 cells). Given the wide range of sizes in physiologically relevant spheroids^24^, we opted for analyzing the smaller spheroids formed at 5-7 d. These differences notwithstanding, SKOV3.ip1 spheroids showed little-to-no disaggregation on 3 kPa gels, but showed extensive and near-complete disaggregation on 25 kPa and 125 kPa gels (Figure 11J and K). These results clearly demonstrate that the positive correlation between matrix stiffness and EOC cell morphology, migration, tension, and spheroid disaggregation is maintained even in highly metastatic cells.

**Figure 11.**
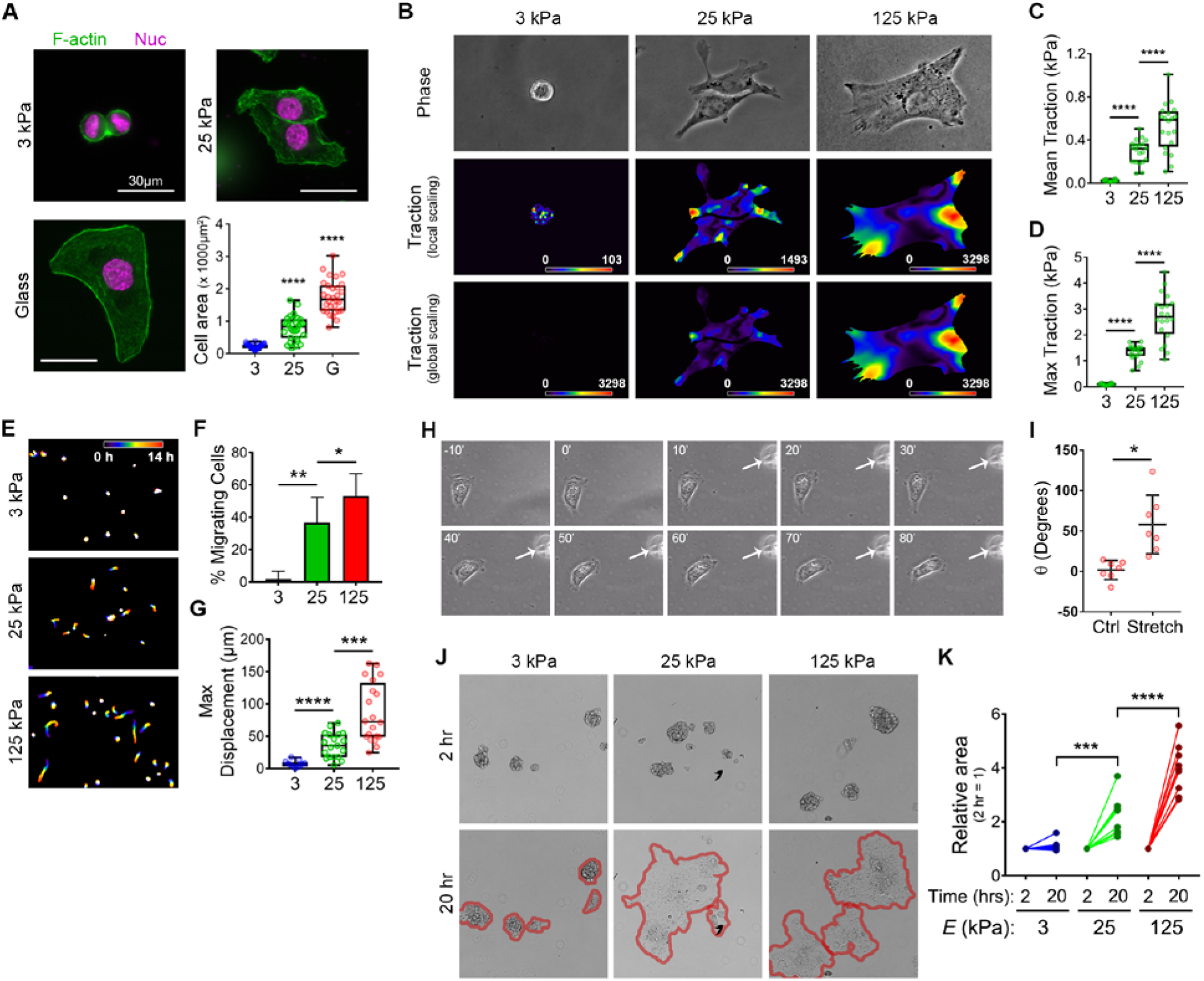
Substrate rigidity positively regulates cell morphology, migration, traction, and spheroid disaggregation in highly metastatic SKOV3.ip1 cells. SKOV3.ip1 cells were cultured to analyze the following experimental parameters: (A) Cell morphology, as described in Figure 2 (graph depicts all data points (colored symbols), with box & whisker plots as described earlier; n = 30 cells from 3 experiments for all samples; ****, *p* < 0.0001); (B-D) Traction force, as described in Figure 4 (n = 20 cells from 3 experiments per condition); ****, *p* < 0.001); (E-G) Cell migration, as described in Figure 5 (panel (E) shows temporal color-coded tracks of the positions of cell nuclei over time; panel (F) shows mean percent migrating cells (± s.d.; n = 80 cells from 3 experiments; *, *p* = 0.0287; **, *p* = 0.0048); panel (G) shows maximum displacements as box & whisker plots as described earlier (n = 11, 19, and 22 cells from 2 experiments for 3, 25, and 125 kPa samples, respectively; ***, *p* = 0.001; ****, *p* < 0.001)); (H, I) Durotaxis, as described in Figure 6 (white numbers indicate time in minutes before or after the hydrogel was stretched in the direction indicated by the arrow; n = 7 cells each condition; *, *p* = 0.002); (J, K) Spheroid disaggregation, as described in Figure 7 (because of the heterogeneous size of SKOV3.ip1 spheroids, the graph in (K) depicts the *relative* area of each spheroid measured at 2 h and 20 h; n = 10 spheroids for each condition; ***, *p* = 0.007; ****, *p* < 0.001).

## Discussion

The current work establishes EOC as a mechano-responsive malignancy. Using polymer hydrogels that mimic the Young’s modulus values found in normal peritoneum – an organ targeted with grave effect during EOC metastasis – we have shown that increasing matrix stiffness enhances a variety of cellular processes germane to EOC dissemination and invasion, including EOC cell morphology, migration, tension, YAP1 nuclear import, and spheroid disaggregation.

In contrast to other epithelial tumors, ovarian cancers predominantly metastasize by sloughing away from primary tumors and moving throughout the abdominal cavity in ascites fluid with the normal peritoneal fluid flow^20^. The mesothelium and the basement membrane beneath the mesothelial cell layer contain numerous ligands that support adhesion and migration such as fibronectin, laminin, type IV collagen and mesothelin. Spheroid disaggregation has been observed on a wide variety of ECM proteins^62^ and the generation of spheroid aggregates requires cellular contractility and correlates with the contractile nature and invasive phenotype of the EOC itself^27,62,90^. Iwanicki *et al* elegantly demonstrated that EOC spheroids exert myosin-dependent mechanical force on underlying mesothelial cells, leading to disruption and displacement of the mesothelial monolayer and providing the EOC cells access to the underlying matrix^25^. The current work suggests, however, that even after the mesothelial barrier is breached, the mechanical properties of the underlying extracellular matrix may contribute to EOC spheroid disaggregation and local invasion.

It is important to note that the elastic modulus values reported and used herein to model and manipulate the mechanical microenvironment were based on analysis of murine peritoneum. Indeed, the current results justify the effort to extend our analysis to normal – and perhaps pathologically altered – human peritoneum. That caveat notwithstanding, it is interesting to note that neither of the EOC cell lines used here showed appreciable spheroid disaggregation on gels with a stiffness equivalent to that of the average value of the peritoneum. This suggests that, *in*vivo, EOC spheroids may either ‘roll’ on the peritoneal surface, sampling the mechanical microenvironment for a region stiff enough to support stronger adhesion and disaggregation or alternatively, after adhesion to a soft region, may elicit a desmoplastic response that leads to stiffening *in situ* and thereby supports disaggregation and, ultimately, localized invasion. This latter hypothesis is comparable to the situation in breast cancer and other tissues^9,10,15,57^ wherein there is a ‘dialog’ between a tumor and its associated stroma that increases the rigidity of the microenvironment, which in turn enhances local invasion.

Indeed, the role that EOC cells have in shaping the mechanical properties of the sites of metastases remains to be investigated. The effect of tumor cells on remodeling the ECM has been extensively studied in breast cancer where matrix metalloproteinases (MMPs) degrade the extracellular matrix and matrix reorganizing proteins such as lysyl oxidases (LOXs) increase collagen cross-linking and, thus, increase tissue tensile strength^13^. The stiffened ECM actively signals to promote integrin-dependent focal adhesions and enhance the activity of pro-migratory signaling pathways such as PI3K^10^. We have previously shown that SKOV-3 cells require MMP activity to degrade the ECM during invasion through matrigel and that leading edge protein kinase A signaling is required for SKOV-3 cell migration and invasion^91^. Other studies have shown that the expression of ECM remodeling proteins such as LOX1 and TGT2 are increased in highly aggressive ovarian cancers^26^. Given the fact that EOC cells sense, respond, and can contribute to the mechanical properties of the ECM, it is tempting to hypothesize a mode of metastasis in which, after dissemination, EOC spheroids not only attach to and clear the mesothelium, but also instigate a program of ECM remodeling characterized by MMP-mediated matrix degradation, LOX1/TG2-mediated matrix cross-linking, and consequent or parallel initiation of a localized inflammatory response^17,19,20^. To this point, there is considerable evidence highlighting the importance of EOC-driven cancer-associated fibroblasts and other aspects of EOC-associated reactive stroma in disease progression and severity^92–102^. Thus, these desmoplastic events would be expected to cause an increase in ECM stiffness which would, in turn, activate integrin signaling, promote focal adhesion formation, and activate migration-associated signaling pathways to ultimately reinforce the locally invasive behavior of EOC cells. Current efforts are underway to assess the ability of EOC spheroids to induce mechanical changes in the peritoneal microenvironment and the contribution of such changes to the local invasion – and ultimately the widespread dissemination – of EOC *in vivo*.

Finally, the results reported here are in stark contrast to those from an earlier effort examining the relationship between matrix stiffness and ovarian cancer cell behavior, which reported that soft, rather than stiff, substrates preferentially supported increased EOC cell adhesion, migration, phospho-MLC intensity, and the magnitude and polarity of traction forces^89^. This is despite the use of both the same cell line (SKOV3) and nearly the same values for substrate rigidity (~3 kPa & ~30 kPa) for the majority of both studies. That report also observed that a pliable microenvironment promoted increased expression of markers of epithelial-mesenchymal transition (e.g. vimentin, N-cadherin) in SKOV3 cells. While the current work did not directly assess these canonical EMT markers, our data showing increased nuclear localization of YAP1 on stiffer matrices suggest a contrary, positive correlation between the microenvironmental rigidity and EOC aggression at the transcriptional level. Finally, the previous report also suggested that this inverse mechanical relationship was a function of metastatic potential, as a less aggressive cell line of different origin (*i.e*. OVCAR3 cells) did not show these behaviors^89^. This, again, is at odds with the current observations that SKOV3 cells SKOV3.ip1 cells, a more aggressive and metastatic subline derived directly from SKOV3 cells, maintained a directly proportional response to matrix rigidity.

The reasons underlying the opposing observations in these two reports are not known. As stated, both studies used the same cell line and hydrogels of comparable mechanical properties for the bulk of the work. One salient difference might be the culture conditions, as the previous work reported using a minimal medium (Hank’s balanced salt solution) without serum for most assays^89^, while the current work used serum-containing medium. We chose our culture conditions because of the biochemical complexity of ascites fluid and, from this, the high likelihood that EOC cells and/or spheroids would be constantly exposed to a rich, heterogeneous mixture of bioactive factors, which is routinely approximated *in vitro* by complex media containing animal sera. This technical difference is noteworthy, as the earlier report showed that some rigidity-dependence of some experimental endpoints (*i.e*. migration, aspect ratio, traction stress, and traction polarization) was either reversed or normalized by treatment with lysophosphatidic acid (LPA;^89^), an abundant component of both serum and malignant ascites that promotes migration and contractility principally through activation of Rho-dependent signaling pathways^103,104^.Another noteworthy difference between the methodology of these two studies is the use of different ECM proteins; specifically, collagen I^89^ and fibronectin (this study). While the use of fibronectin is relevant and appropriate (see *Results*, above), both proteins – as well as several other ECM proteins and components – are assuredly physiologically relevant to EOC progression and dissemination, albeit quite likely at different stages of these processes^18,40^. Importantly, it is becoming increasingly evident that different matrix proteins, acting through different integrins and associated adhesive contacts, can elicit distinct profiles of mechanosensitive behavior^68,105-107^, although the true physiological relevance of these differences is not yet fully understood. It would be intriguing and informative to directly assess whether EOC cellular mechano-response can effectively reverse itself based on matrix- and/or integrin-specific inputs; efforts along these lines are underway. Nonetheless, the results reported herein unequivocally demonstrate that increasing mechanical stiffness in the microenvironment can enhance EOC cell adhesion, migration, and spheroid disaggregation, and suggest that this relationship may contribute to EOC invasion and disease progression.

## Methods

### General cell culture

The human epithelial ovarian cancer cell line SKOV3 (American Type Culture Collection) and its more invasive and metastatic sub-clone SKOV3.ip1 (a kind gift from D. Schlaepfer, UCSD) were maintained in a humidified incubator at 37°C containing 5% CO2 in DMEM supplemented with 10% fetal bovine serum. Cells were trypsinized and split 1:5 every 3-4 days to maintain at subconfluency. Both cell lines were authenticated approximately every 4 months by the University of Vermont Advanced Genome Technologies Core Facility, using the GenePrint 10 short tandem repeat analysis system (Promega Corporation).

### Antibodies and miscellaneous reagents

Antibodies (and their source) against the following targets were used: paxillin (Abcam, ab32084); phospho-myosin light chain 2 (pThr18/pSer19; Cell Signaling Technology, #3674); myosin light chain 2 (Cell Signaling Technology, #3672); vinculin (hVin-1; Millipore-Sigma V9131); phospho-FAK (pTyr397; Cell Signaling Technology, #3283); FAK (C-20; Santa Cruz Biotechnology; sc-558); YAP1 (D8H1X; Cell Signaling Technology, #14074). DAPI (4’,6-diamidino-2-phenylindole dihydrochloride), Hoechst 33342, and phalloidins conjugated to Alexa Fluor 488, 594, or 647 were from ThermoFisher Scientific. Fasudil (also known as HA-1077) was from Cayman Chemical, PF-562271 was from ChemScene, and Blebbistatin (#203389) and latrunculin A (L5163) were from Millipore Sigma. Acrylamide and N,N-methylenebisacrylamide were purchased from National Diagnostics. Human plasma-derived fibronectin (Corning #356008) was from Thermo Fisher. Glass-bottom imaging dishes were from Cellvis. Other chemicals and reagents, unless otherwise noted, were purchased from Millipore Sigma.

### Image analysis and figure preparation

Unless otherwise stated, images were adjusted and analyzed with the Fiji distribution of ImageJ^108,109^, using built-in functions, plugins, or custom macros as indicated. All graphs were generated and statistics analyzed using GraphPad Prism. Figures were made using Photoshop Creative Cloud.

### Fabrication of polyacrylamide hydrogels

Polymer hydrogels were made essentially as previously described^110^. After sterilizing with 70% ethanol, 22mm coverslips or glass bottom dishes (Cellvis) were treated with 2 N NaOH for 15 min and with 3-aminopropyltriethoxysilane (APTMS; Millipore Sigma) for 3 min. Coverslips were washed 3 x 5min with ddH_2_O then incubated with 0.5% glutaraldehyde for 30 min. Activated coverslips were either used immediately or stored desiccated for up to 2 weeks. Polyacrylamide gels contained final concentrations of 7.5% or 12% acrylamide and either 0.5% or 0.05% bis-acrylamide. These concentrations were chosen based on published values^33^ to give desired Young’s elastic modulus values, which were then confirmed by atomic force microscopy as described below. Polymerization was initiated with 2.5 μl 10% APS and 0.5 μl TEMED (N,N,N’,N’-tetramethyl-ethylenediamine). Gels were cast onto activated coverslips or glass bottomed dishes by adding a 25μl drop of the gel solution onto the activated surface and overlaying another 22 mm coverslip passivated with Rain-X (to facilitate its removal after polymerization), yielding a gel height of approximately 65 μm. Gels were allowed to polymerize for 30 minutes and, after removal of the top coverslip, were washed 3 x 5 min in 50 mM HEPES pH 8.5, then incubated with 0.4 mM sulfosuccinimidyl 6-(4’-azido-2’-nitrophenylamino)hexanoate (sulfo-SANPAH; Thermo Fisher) for 90 seconds at room temperature 4” from a 364 nm UV light source (IntelliRay 400; Uvitron). Activated gels were washed 3 x 5 min in 50 mM HEPES pH 8.5 and incubated with 20 μg/ml fibronectin (FN) diluted in 50 mM HEPES pH 8.5 for 1 h at 37°C. FN-coated gels were sterilized for 15min under UV light, washed (3 x 5 min) in PBS and either used immediately or stored at 4°C for up to 1 week.

### Obtaining and mounting peritoneal tissue

All mouse tissue was obtained following standard IACUC protocols for tissue sharing. All mice were 6-10 week-old female C57BL6 devoid of any ostensible fibrotic, abdominal, or peritoneal pathologies. Once sacrificed by CO2 inhalation and cervical dislocation, mice were skinned and 1 x 2 cm sections of abdominal peritoneum were dissected and adhered to glass slides visceral side-up using Cell-Tak (BD Biosciences), maintaining the pre-dissection dimensions of the tissue. A pap-pen was used to create a hydrophobic ring around the tissue and the circumscribed tissue was covered with PBS for imaging and mapping.

### Microindentation atomic force microscopy and force mapping

Force indentation curves were obtained from peritoneal tissue or hydrogels using a 5.0 μm borosilicate glass sphere probe on a silicon nitride cantilever (spring constant: 0.06 N/m; NovaScan) using the contact mode on a MFP-3D BIO Atomic Force Microscope controlled by an IGOR Pro interface (Asylum Research). Probe spring constants were confirmed using thermal oscillations in air before each imaging session. Deflection curves were obtained using the force map as follows; a 20 μm x 20 μm square sample was divided into an 8 x 8 grid, yielding 64 2.5 μm x 2.5 μm grid-squares and a single deflection curve for each. The probe speed was set to 10 μm/s and retracted 5 μm between each point to prevent adhesion of the sample to the probe. At least 3 random 20 μm x 20 μm force maps were obtained on each sample, generating at least 192 points for each sample. Care was taken to avoid any unadhered portions at the edges of tissues.

Young’s modulus for each deflection curve was calculated by fitting the deflection curve to the Hertz model in the IGOR software package. Values below 15% and above 80% of the curve were excluded to eliminate potential artifacts. Once the Young’s modulus was determined for each deflection curve in a force map, the values were imported into a custom-written ImageJ macro to generate color-coded heat maps depicting relative values within the grid squares. All statistical analyses were done with GraphPad Prism software.

### Immunofluorescence, morphometry, and co-localization analyses

Glass coverslips (22 or 25 mm round; Fisher Scientific) were sterilized by sequential 30 min soaks in 70%, 95%, and 100% EtOH and then air-dried in a tissue culture hood. Cells were fixed in 3.7% formaldehyde in Tris-buffered saline (TBS) for 10 min, permeabilized for 10 min in TBS containing 0.25% triton X100, blocked with TBS containing 5% BSA for 1 h at RT then incubated with antibodies against paxillin (1:500), phospho-MLC2 (1:500), vinculin (1:400), or phospho-FAK (1:250) for 1 h at RT. After washing with TBS, cells were incubated with Alexa-647 coupled donkey anti-mouse, Alexa-596 coupled donkey anti-rabbit secondary antibodies (Invitrogen, 1:400) and Alexa-488 conjugated phalloidin (Invitrogen, 1:100) for 1 h at RT. Coverslips were washed then mounted onto slides using PermaFluor (ThermoFisher). Epifluorescence images were captured though a 60X Plan Apo oil immersion objective on a Nikon Eclipse TE2000E or TiE inverted microscope using the appropriate fluorophore-specific filters (Chroma Technology Corp) and a CoolSnap HQ camera (Photometrics) controlled by Nikon Elements software. Images were analyzed in ImageJ. For cell area, a binary mask was created by thresholding phalloidin images and area was determined using the ImageJ *Measure* tool.

For immunofluorescence of YAP1, cells were plated at a density of 55,000 cells per 35 mm well on 3kPa, 25kPa, 125kPa and glass substrates overnight at 37°C in DMEM containing 10% FBS. Cells were fixed 18hr post plating for 15 minutes in 4% paraformaldehyde in PBS, and blocked and permeablized in 5% donkey serum + 0.3% triton X-100 in PBS for 1hr at room temperature, then incubated with anti-YAP1 antibody (1:100) in 1% BSA and 0.3% triton X-100 in PBS overnight at 4°C. After washing with PBS, cells were incubated with Alexa-488 coupled donkey anti-rabbit (Abcam, 1:400) and Alexa-594 conjugated phalloidin (Molecular Probes, 1:100) for 90min at room temperature. Coverslips were washed with 5 times for 5min each with PBS, followed by a 5 min incubation with DAPI (1:6000) in PBS. Coverslips were mounted onto slides using PermaFluor (Thermofisher) and epifluorescent images were captured through a 40x Plan Fluor oil immersion objective.

For assessment of morphological effects of mechanotransduction inhibitors, cells plated on fibronectin-coated 25 kPa hydrogels were allowed to adhere overnight and then treated with DMSO (0.1%), blebbistatin (25 μM), Fasudil (10 μM), or PF-562271 (0.25 μM) for 30 minutes. Cells were fixed in 3.7% formaldehyde for 20 minutes, permeabilized for 5 minutes in 0.25% triton X100, blocked in TBS containing 3% BSA for 1 hour at room temperature, then incubated with antibodies against phospho-MLC (1:200), vinculin (1:400), or phosphor-FAK (1:100) for 1 hour at room temperature. After washing with TBS, cells were incubated with Alexa-488 coupled donkey anti-mouse, Alexa-488 coupled donkey ani-rabbit, or Alexa-594 coupled donkey antirabbit secondary antibodies (1:400; Abcam), and Alexa-594 conjugated phalloidin (1:100; Invitrogen) for 1 hour at room temperature, then washed, mounted, and imaged as described above.

### Immunoblotting

Cells on polyacrylamide gels or glass coverslips were lysed by inversion onto a 100 μl drop of Triton X-100 lysis buffer (1% Triton X-100; 50 mM Tris pH 7.2; 10% glycerol; 25 mM ß-glycerophosphate; 2 mM EDTA; 2 mM EGTA) containing protease and phosphatase inhibitors and incubation on ice for 10 min. Lysates were gently collected, incubated on ice an additional 10 min, then cleared by centrifugation and assayed for protein content (BCA assay; ThermoFisher). Samples were mixed with 5x Laemmli sample buffer, boiled, separated on 15% SDS-PAGE gels, and transferred to PVDF membranes. After blocking for 1 h (in 1.5% gelatin in TBS-T), membranes were incubated with antibodies against phosphor-MLC2 (1:500) or total MLC2 (1:500) overnight at 4°C. The membranes were washed in TBS-T, incubated with appropriate HRP-conjugated secondary antibodies (1:5000; Calbiochem) for 60 min, washed again, then developed by enhanced chemiluminescence (Pierce ECL Plus; ThermoFisher).

### Traction force microscopy

Polyacrylamide hydrogels for traction force microscopy were prepared as previously described (Knoll et al., 2014). Briefly, acid washed glass coverslips (18 mm) were incubated in poly-L-lysine (10 μg/mL in sterile water) for 1 hour at room temperature, then coated with 0.2 μm carboxylate-modified red FluoSpheres (1:200 in sterile water; Thermo Fisher) for 15 minutes at room temperature. Bead coated coverslips were used as top coverslips to cast polyacrylamide hydrogels on functionalized, glass bottom imaging dishes as described above. Hydrogels were allowed to polymerize for 30 minutes at room temperature and washed for 30 minutes in 50 mM HEPES (pH 8.5) before the top coverslips were removed. Beaded gels were then washed again in 50 mM HEPES and coated with fibronectin (20 μg/mL in HEPES pH 8.5) for 1 hour at 37°C. Cells were allowed to adhere overnight and transferred into CO2 independent medium (Thermo Fisher) supplemented with 10% fetal bovine serum prior to imaging. Phase contrast and fluorescent bead images were captured through a 20x or 40x Plan Apo objective on a Nikon Eclipse TE-2000E inverted microscope. Spheroids were seeded onto beaded gels as described above and imaged 2 h and 20 h after seeding. Bead positions were imaged at the indicated times after cell/spheroid seeding, as well as after clearing cells or spheroids by the addition of 0.3% SDS (to obtain a zero-force image).

Bead images were assembled into stacks and registered using the MultiStackReg plugin for ImageJ. The resulting registration transformation matrix was then applied to the corresponding phase images. Traction force microscopy was performed using the PIV and FTTC ImageJ plugins (created and generously provided to the research community by Q. Tseng and available at https://sites.google.com/site/qingzongtseng/imagejplugins), called by a custom-written macro to process stacks. Typically, three passes of advanced iterative PIV were performed with first-pass parameters as follows: interrogation window size = 128; search window size = 256; vector spacing = 64. Subsequent passes stepped those parameters down by half and the correlation threshold was set at 0.8. Fourier transform traction cytometry (FTTC) was performed using a Poisson ratio of 0.5, pixel size of 0.161 microns, and a Young’smodulus value appropriate for the hydrogel being analyzed. Using an L-series sweep as described elsewhere^111,112^, we found that optimal regularization factor values varied as a function of hydrogel rigidity; thus, we used regularization factors of 2.5 x 10^−10^, 2 x 10^−10^, and 1 x 10^−10^ for 125 kPa, 25 kPa, and 3 kPa gels, respectively. Cell contours in phase contrast images were outlined manually and transferred to traction force maps; values outside the contours were cleared to depict only forces within the area of the cells themselves. Maximum and mean traction forces were measured from the 32-bit force maps, while total cell strength was calculated as the product of the cell area (in μm^2^) and the mean traction force (in kPa).

### Live cell imaging, cell tracking, and migration analyses

Cells plated on hydrogels or glass-bottom dishes were incubated overnight in DMEM with 10% FBS. For cell tracking, 3.5 x 10^4^ cells were seeded in the dishes and incubated overnight. The nuclei were then stained with Hoescht 33342 (1:2000 in Ringer’s buffered saline (10 mM HEPES; 10 mM glucose; 155 mM NaCl; 5 mM KCl; 2 mM CaCh; 1 mM MgCh)) for 10 min at room temperature followed by washing twice with Ringer’s saline. The cells were re-fed with DMEM with 10% FBS and monitored by live cell microscopy using a Lionheart FX automated live cell imager (Biotek) with a 10x dry objective (UPLFLN; Olympus), maintaining the cells at 37° C in a 5% CO2 environment for the duration of the imaging. Phase contrast and fluorescent images were captured every 15 min for 14 h. The nuclei tracking recipe of SVCell/AIVIA (DRVision Technologies, Bellvue, WA) was used to track individual nuclei and obtain (x,y) coordinates to calculate displacement, path length, and cell speed. Migrating cells were identified as those that migrated beyond a circular area 2x the diameter of the cell over the 14 h course of imaging. Maximum displacement calculated as the largest change in Euclidean distance of a given cell over the entire course of imaging.

### Durotaxis assays

Cells were seeded, mounted, and maintained on the microscope as above and were manipulated with a glass microneedle as described previously^113^. Micropipettes were fashioned from borosilicate glass capillaries (World Precision Instruments) on a two-stage Pul-2 pipette puller (World Precision Instruments). A Narishige MF-900 microforge was used to form the micropipette tip into a hook with a rounded end to engage the polyacrylamide hydrogels without tearing. The forged microneedle was mounted on a micro-manipulator (Leica Leitz mechanical or Narishige MHW-3) and lowered onto the gel surface approximately 20 μm away from a cell. Once engaged, the gel was pulled approximately 20 μm in a direction perpendicular to the cell’s axis of migration. Quantification of durotactic response was calculated as the angle defined by the long axis of the cell immediately prior to durotactic stretch and its long axis 75 min after stretch, and was compared to the long axis angles of unstretched motile cells at 0 and 75 minutes.

### Spheroid disaggregation assay

Multi-cellular spheroids of SKOV3 and SKOV3.ip1 cells were generated as follows: Cells were trypsinized, collected in Medium 199 (Thermo Fisher) + 10% FBS (*n.b*. the use of Medium 199 as opposed to DMEM was found to significantly enhance the formation and health of spheroids from both lines), and diluted to 20,000 cells / ml in the same medium. The wells of a 96-well plate were coated with 0.1 ml of 1% agarose (UltraPure; Millipore Sigma) in Medium 199, allowed to cool to room temperature, then seeded with 0.1 ml of the cell suspension (*i.e*. 2000 cells/well). After incubation under standard culture conditions, SKOV3 cells condensed into single, compact spheroids with ~100% efficiency within 3 d, while SKOV3.ip1 aggregated into smaller, more numerous spheroids over 5-7 d. Spheroids were collected by gentle trituration with a short glass Pasteur pipette and collected into 1 ml M199/10% FBS. The spheroids were pelleted at 100x g for 5 minutes, 95% of the supernatant medium was removed and replaced with 6 mls fresh medium, and the spheroids were gently resuspended and dispersed evenly into six-well plates containing hydrogel-coated coverslips. Spheroids were allowed to attach for 2 h before phase contrast imaging with 10x or 20x Plan Apo objectives. Spheroids were then incubated overnight and images were taken 18 h after the initial images. To quantify the degree of dispersion, images were thresholded using ImageJ to outline the periphery of the aggregate, and total area of the 20 h image was divided by the area of the initial image. Where indicated, spheroids were seeded onto hydrogels containing fluorescent nanospheres and random fields at the periphery of the spheroids were imaged after 4 h for assessment of cellular traction forces as described above.

### Data availability

The datasets generated during and/or analyzed during the current study are available from the corresponding author on reasonable request.

## Acknowledgements

The authors thank David Schlaepfer (U.C.S.D) for providing SKOV3.ip1 cells, Jason Stumpff (UVM) for helpful discussions, and Jonathan Patterson for technical assistance. This work was supported by NIH grants R01GM097495 and R01GM 117490 (to AKH) and a University of Vermont Cancer Center (UVM) and Norris Cotton Cancer Center (Dartmouth) Collaborative Research Program grant (to AKH).

## Author contributions

A.J.M. helped conceive and plan the project, performed experiments and analyzed data to assess cell morphology, immunofluorescence patterns, cell migration, durotaxis, and spheroid disaggregation, and helped write the manuscript. S.R.H. performed all of the AFM experiments as well as experiments to assess cell migration. K.V.S. performed experiments and analyzed data to assess durotaxis and YAP1 localization and helped prepare figures and write the manuscript. H.N. performed experiments and analyzed data to assess cell morphology, immunofluorescence patterns, migration, and traction force, and helped write the manuscript. Z.L.E. performed experiments and analyzed data to assess cell morphology, migration, and YAP1 localization. A.K.H. conceived and planned the project, trained all personnel involved, procured funding, performed experiments and analyzed data, and wrote the majority of the manuscript.

---

**Supplemental Figure 1.**
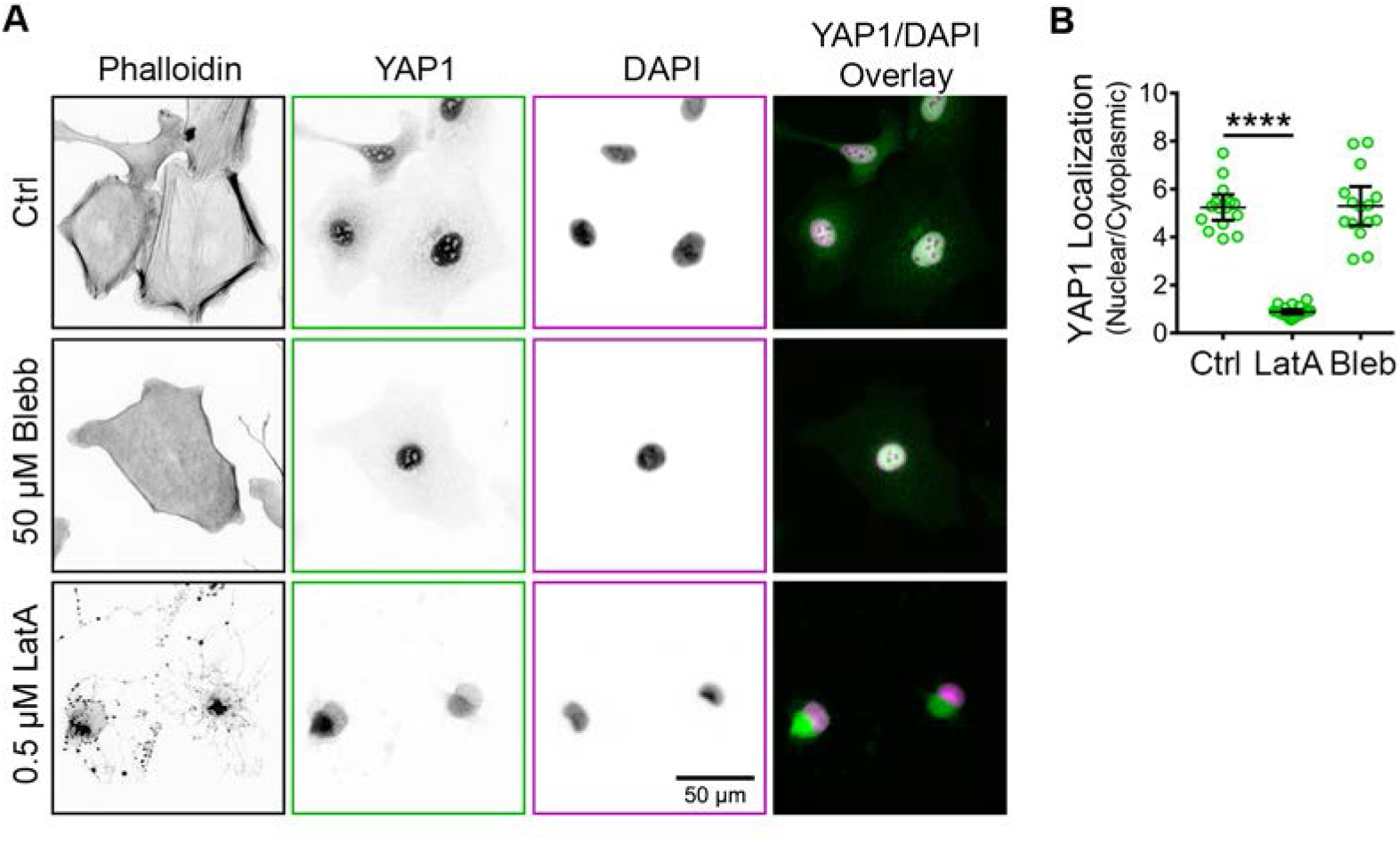
Nuclear translocation of YAP1 is independent of actomyosin contractility but dependent on F-actin cytoskeleton integrity. (A) YAP1 localization shown for representative cells exposed to X μM DMSO (ctrl, top row), 50 μM blebbistatin (Blebb), or 0.5 μM latrunculin A (Lat A) for 4 hours before being fixed and stained to visualize F-actin, YAP1, and nuclei. (B) Quantification of YAP1 localization given as a ratio of YAP1 signal in the nucleus to YAP1 signal in the cytoplasm for cells in each treatment condition. The graph depicts all measure values (colored symbols) and the mean values (± s.d.; n = 15 for DMSO and latrunculin A and 21 for blebbistatin; *p* = < 0.0001 using the Mann-Whitney test).

